# Impacts of Deep-Sea Mining on Microbial Ecosystem Services

**DOI:** 10.1101/463992

**Authors:** Beth N. Orcutt, James Bradley, William J. Brazelton, Emily R. Estes, Jacqueline M. Goordial, Julie A. Huber, Rose M. Jones, Nagissa Mahmoudi, Jeffrey J. Marlow, Sheryl Murdock, Maria Pachiadaki

**Affiliations:** Bigelow Laboratory for Ocean Sciences, 60 Bigelow Drive, East Boothbay, ME, 04544, USA; University of Southern California, Zumberge Hall of Science, Los Angeles, CA, 90089, USA; University of Utah, School of Biological Sciences, 257 South 1400 East, Salt Lake City, UT, 84112, USA; University of Delaware, College of Earth, Ocean & Environment, 700 Pilottown Road, Lewes, DE, 19958, USA; Woods Hole Oceanographic Institution, 266 Woods Hole Road, Woods Hole, MA, 02543, USA; Harvard University, Department of Organismic & Evolutionary Biology, 16 Divinity Ave, Cambridge, MA, 02138, USA; McGill University, Department of Earth & Planetary Science, 3450 University Street, Montreal, QC, H3A 0E8, Canada; University of Victoria, School of Earth and Ocean Sciences, 3800 Finnerty Road, Victoria, BC, V8W 2Y2, CanadaCanada

**Author notes:** **Corresponding author:** Beth N. Orcutt, Bigelow Laboratory for Ocean Sciences, 60 Bigelow Drive, East Boothbay, ME, 04544, USA.

**Keywords:** deep-sea mining, ecosystem services, hydrothermal vents, inactive sulfides, ferromanganese nodules, cobalt crusts, seamounts, marine microbiology

## Abstract

Interest in extracting mineral resources from the seafloor through deep-sea mining has accelerated substantially in the past decade, driven by increasing consumer demand for various metals like copper, zinc, manganese, cobalt and rare earth elements. While there are many on-going discussions and studies evaluating potential environmental impacts of deep-sea mining activities, these focus primarily on impacts to animal biodiversity. The microscopic spectrum of life on the seafloor and the services that this microbial realm provides in the deep sea are rarely considered explicitly. In April 2018, a community of scientists met to define the microbial ecosystem services that should be considered when assessing potential impacts of deep-sea mining, and to provide recommendations for how to evaluate these services. Here we show that the potential impacts of mining on microbial ecosystem services in the deep sea vary substantially, from minimal expected impact to complete loss of services that cannot be remedied by protected area offsets. We conclude by recommending that certain types of ecosystems should be “off limits” until initial characterizations can be performed, and that baseline assessments of microbial diversity, biomass, and biogeochemical function need to be considered in environmental impact assessments of all potential instances of deep-sea mining.

## 1. INTRODUCTION

With increasing demand for rare and critical metals – such as cobalt, copper, manganese, tellurium, and zinc – there is increasing interest in mining these resources from the seafloor (Hein et al., 2013a; Wedding et al., 2015). The primary mineral resources in the deep sea that attract attention fall into four categories (Figures 1, 2): (1) massive sulfide deposits created at active high-temperature hydrothermal vent systems along mid-ocean ridges, back-arc spreading centers, and volcanic arcs; (2) similar deposits at inactive hydrothermal vent sites; (3) polymetallic “nodules” that form on the seafloor of the open ocean (often referred to as manganese nodules); and (4) other polymetallic crusts that can form in the deep sea at underwater mountains called seamounts (often referred to as cobalt crusts). Current areal estimates of these resources range from 38 million km^2^ for ferromanganese nodules, 3.2 million km^2^ for massive sulfides (combined active and inactive), and 1.7 million km^2^ for polymetallic crusts on seamounts (Petersen et al., 2016). Some of these resources occur within the Exclusive Economic Zones (EEZ) of coastal nations, while others occur in international waters. In some EEZs continental shelf sediments, additional commercial interests include diamond and phosphorite deposits. Since these fall exclusively within national jurisdictions, they will not be a focus of this article, although mining activities for these deposits are currently occurring (Miller et al., 2018).

**Figure 1.**
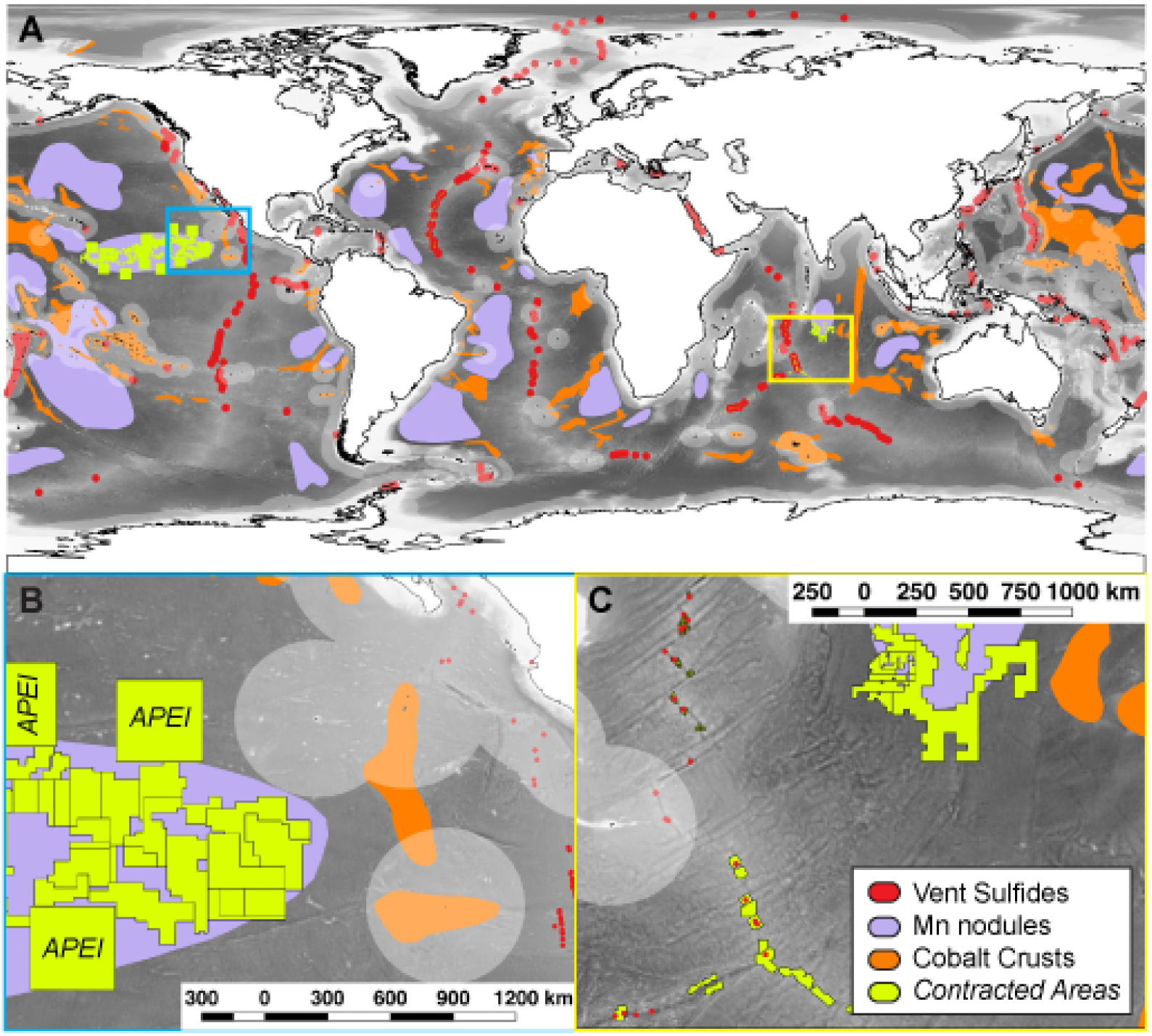
Locations of deep-sea mineral resources and current exploration contract zones. In all panels, areas of the seafloor within nations’ Exclusive Economic Zones (EEZ) highlighted in light grey boundaries along coastlines, whereas remaining seafloor within “the Area” not shaded. Panel A – Global seafloor distribution of hydrothermal vent (active and inactive) polymetallic sulfide deposits (red), ferromanganese nodules (purple), and cobalt crusts on seamounts (orange) overlain by current exploration contract zones (green) issued by the International Seabed Authority (ISA). Panel B – Highlight of exploration contracts and Areas of Particular Ecological Interest (APEI) in the eastern region of the Clarion Clipperton Zone and the East Pacific Rise vent locations on the western edge of Mexico, as shown by the blue bounding box in A. Panel C – Highlight of vent sites and contracted zones along part of the Southwest Indian Ridge and nodule exploration contracts in the Indian Ocean, as shown by the yellow bounding box in A. Underlying maps generated with data from the U.S. National Oceanographic and Atmospheric Administration (NOAA; coastlines), the General Bathymetric Chart of the Oceans (GEBCO) hosted by the British Oceanographic Data Centre (gridded bathymetry data), and the marineregions.org database (EEZ). Shape file information for nodules and cobalt crusts from (Hein et al., 2013a), for polymetallic sulfides InterRidge Vents Database version 3.3 hosted by the Institut de Physique du Globe de Paris (credit S. Beaulieu, Woods Hole Oceanographic Institution, 2015, funded by the U.S. National Science Foundation #1202977), and for ISA exploration contract areas from the Deep Sea Mining Watch project version 1.2 hosted by the University of California Santa Barbara Benioff Ocean Initiative.

**Figure 2.**
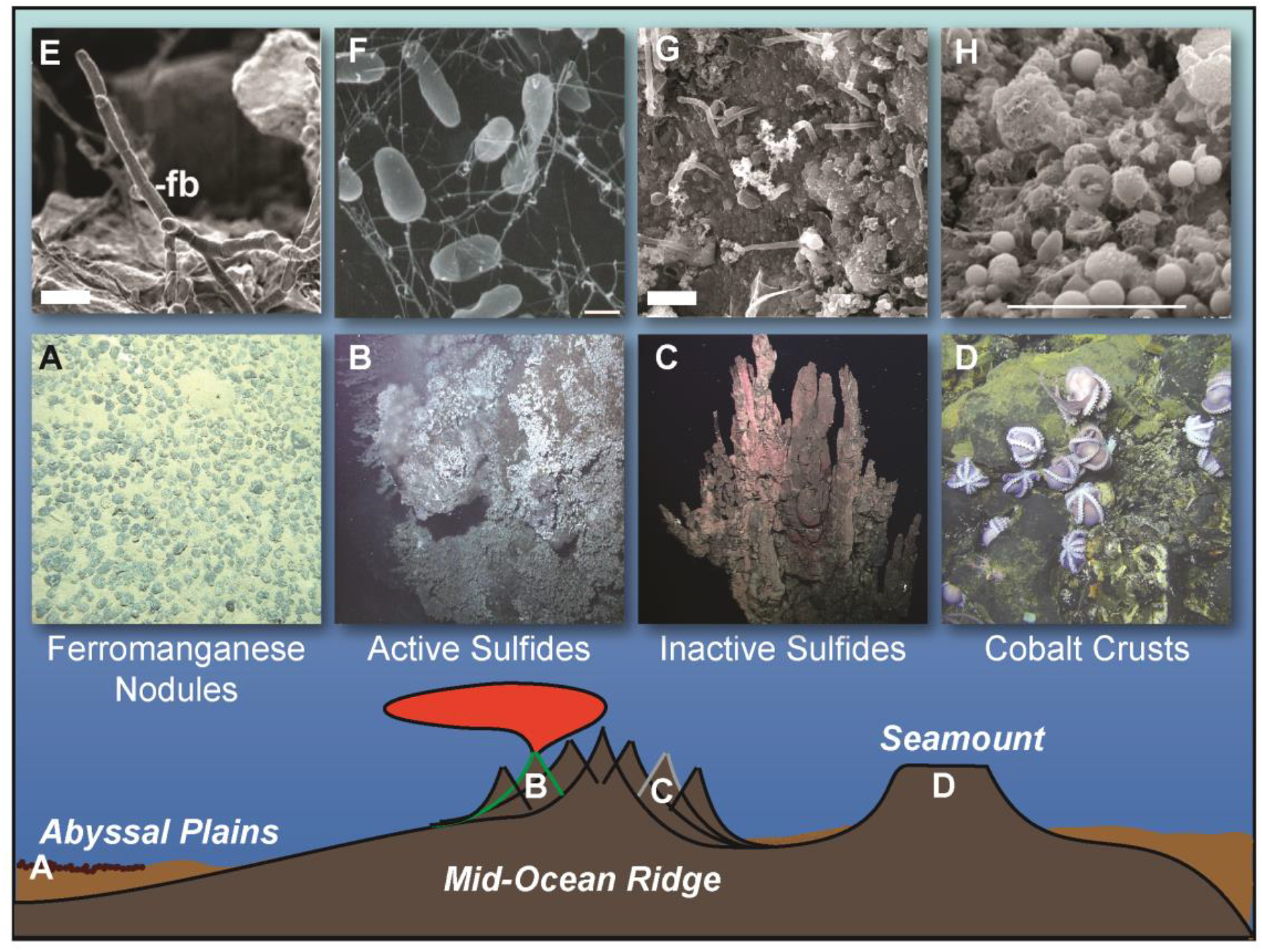
Types of deep-sea habitats with mineable resources. A – Ferromanganese nodules on the sediment of abyssal plains, image modified from Wikipedia courtesy Abramax. B – Active hydrothermal vent sulfide deposits from the Juan de Fuca Ridge, with chemosynthetic animals colonizing the sulfide surface around areas of fluid venting. Photo courtesy of AT15-34 cruise chief scientist Ray Lee, Western Washington University, U.S. National Science Foundation, HOV *Alvin* dive 4420, 2008, © Woods Hole Oceanographic Institution. C – Inactive hydrothermal vent sulfide deposits from the Galapágos Rift. Image by ROV *Hercules* courtesy of NOAA Okeanos Explorer Program 2011 Galapágos Rift Expedition. D – Cobalt-rich crusts that form on seafloor basalts on seamounts, which can be areas of diffusive hydrothermal fluid flow and sites of deep-sea animal brooding. Photo courtesy of AT26-24 Chief Scientist Geoff Wheat, Univ. of Alaska Fairbanks, U.S. National Science Foundation, HOV *Alvin*, 2014, © Woods Hole Oceanographic Institution. Panels E-H show examples of microscopic organisms living on these resources. E – Microbial filaments on a manganese nodule, modified from (Wang et al., 2009); scale bar 1 μm. F – *Desulfurobacterium* bacteria isolated from a deep-sea hydrothermal vent at Axial volcano, courtesy of Julie A. Huber; scale bar 1 μm. G – Rod-shaped cells on inactive sulfides from the East Pacific Rise, modified from (Toner et al., 2013) and courtesy of Brandy Toner; scale bar 2 μm. H – Ferromanganese crust on seafloor basalt from the East Pacific Rise, modified from (Santelli et al., 2008) and courtesy of Cara Santelli; scale bar 10 μm. Cartoon schematic modified from (Schrenk et al., 2009) with permission.

For seabed mineral resources in international waters beyond national jurisdiction (referred to as “the Area”), access is only possible through the International Seabed Authority (the ISA) as established in the United Nations Convention on the Law of the Sea (UNCLOS). The ISS awards contracts of blocks of seafloor via sponsoring States to authorized contractors for resource exploration activity. This is done following regulations established under the Mining Code, which were established in 2000 and updated in 2013 for polymetallic nodules, in 2010 for polymetallic sulfides, and in 2012 for cobalt-rich crusts. More than 1.3 million km^2^ of international seabed is currently set aside in 29 exploration contracts for mineral exploration in the Pacific and Indian Oceans and along the Mid-Atlantic Ridge (Figure 1); another 1 million km^2^ of seabed has been licensed or applied for in waters under national jurisdiction (Cuyvers et al., 2018). Currently, no country has an exploitation license to actively mine these resources in the Area, but the ISA is developing international regulations to govern future exploitation activities within the Area (International Seabed Authority, 2012; 2013). A few States have already begun allowing resource exploration and extraction testing in their national waters (Figure 1). Companies are developing and testing prototype mining equipment for this purpose. Proponents of deep-sea mining argue that resource extraction from the deep-sea is more environmentally friendly than mining on land for the same metals, but the environmental impacts of these efforts are currently poorly understood. The UNCLOS stipulates that deep-sea mining related activities on the international seabed within the Area must be carried out for the benefit of mankind (UNCLOS Articles 136, 137 and 140; (Cuyvers et al., 2018)). Thus, it is imperative to robustly and unbiasedly assess what the positive and negative impacts of deep-sea mining may be, to determine the nature and extent of benefits and consequences for mankind.

Mineral resources on the seabed are also centerpieces of deep-sea ecosystems, functioning as refugia and stepping stones for animal biodiversity. For example, polymetallic crusts in the deep-sea serve as hard substrate for the attachment of sessile animal communities such as sponges, or for egg-laying for mobile species like octopus which do not anchor in the soft sediment surrounding the deposits. As another example, unique animal communities have evolved to survive under the high temperature and extreme chemical conditions found at hydrothermal vents where massive sulfide deposits form from the interaction of these hot, mineral-rich fluids with surrounding cold seawater. Thus, these mineral deposits often host “hotspots” of animal life on the otherwise barren seafloor. There have been several recent studies and syntheses on the potential impacts of mining of these mineral resources on animal life (Boschen et al., 2013; Vanreusel et al., 2016; Jones et al., 2017; Suzuki et al., 2018).

In addition to the visible animal life associated with these resources, diverse *microscopic* life also flourishes in these systems. This nearly invisible microbial life is responsible for the majority of the chemical cycling that occurs in these habitats, providing essential ecosystem services for the deep-sea environments. Microbial life represents a vast and diverse genetic reservoir with mostly unexplored potential for medical and commercial applications. Despite the importance of the microscopic component of life to ecosystem services in the deep sea, this category has been largely overlooked in current planning related to assessing and evaluating possible environmental impacts related to deep-sea mining.

To address this gap in understanding and provide recommendations to policy makers about the possible impacts to deep-sea ecosystem services provided explicitly by microscopic life, a workshop of experts in deep-sea microbial ecology and geochemistry convened in April 2018 to discuss these topics, with support from the Center for Dark Energy Biosphere Investigations, the Deep Carbon Observatory, and the Bigelow Laboratory for Ocean Sciences. The outcomes of this workshop are presented here. We provide an overview of the four mineral resource types along with descriptions of the microbial ecosystems that they support, the ecosystem services that these microbial communities provide, and an assessment of possible impacts to these services from mining activity. We also provide recommendations for baseline assessment and monitoring to evaluate the impact of mining activities on microbial ecosystem services.

## 2. EVALUATING THE POTENTIAL IMPACT OF DEEP-SEA MINING ON ECOSYSTEM SERVICES FROM MICROORGANISMS

Seventy percent of the solid exterior of Earth lies under the ocean. While our perceptions of life on Earth are skewed by our daily encounter with photosynthesis-supported life on land, the deep-sea is a fundamentally different environment where sunlight does not penetrate. In deep-sea environments, energy for life comes in two forms. The first is through the respiration of organic matter delivered either in dissolved or particulate form (ranging from small particles up to large “food falls” like dead whales) from the sunlit surface world that is ultimately sourced from photosynthesis. The second source of energy is through generation of new organic matter (i.e., primary production) from a process known as *chemosynthesis*, where energy from inorganic chemical reactions is used to convert dissolved carbon dioxide into the organic molecules (sugars, fats, proteins, etc.) that are the building blocks of life. The ratio of these two energy sources can vary significantly in the deep sea, with habitats like hydrothermal vents offering a figurative buffet of chemical reactions that can fuel abundant chemosynthesis-driven microbial life. Similarly, in the low-temperature mineral deposits like ferromanganese nodules and cobalt crusts, chemosynthetic processes also occur (Orcutt et al., 2015), although the ratio of the two energy sources is poorly constrained. In these ecosystems, the chemosynthetic microbial life can form the base of the food web. Thus, disruption to the supply of chemical energy sources can have consequences for the amount and type of life that can be supported (Figure 3). The following sections describe how mining activities can upset the chemical energy supplies that fuel microbial life in these ecosystems, and how this can result in a disruption of the ecosystem services that microscopic life provides (Figure 3).

**Figure 3.**
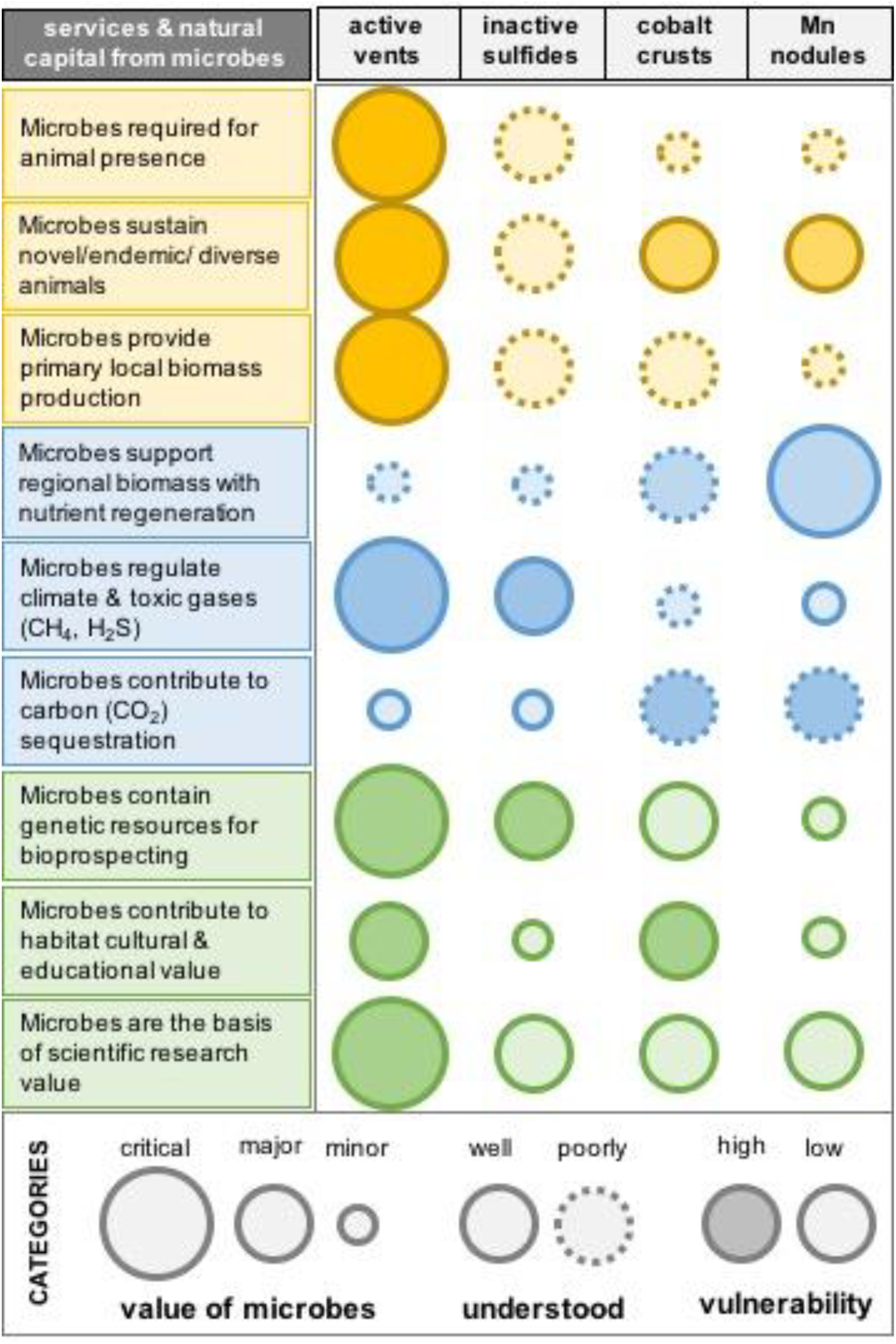
A qualitative assessment of the ecosystem services from microorganisms in deep sea habitats with mineable resources. The size, outline, and shading of symbols reflects the value that microbes support in each system, how well microbial aspects of the ecosystem are understood, and the vulnerability of microbial aspects to mining impacts, respectively, per the legend.

### 2.1. ACTIVE VENTS AND ACTIVE VENT FIELDS

Hydrothermal vents are among the most dynamic environments on Earth, where hot, chemically reduced fluids come into contact with cold, oxidized seawater, leading to the precipitation of metal-rich deposits on and beneath the seafloor surrounding these vents (Figure 2). The high flux of metal-rich fluids mixing with cold, oxic seawater is a natural mechanism for accumulating iron, copper, zinc, and other economically viable elements within metal-sulfide rich mineral deposits, which makes these areas conducive to supporting chemosynthetic life as well as bring attractive targets for mining.

Any given vent ecosystem might only be the size of a football field, with a handful of 5-10 m diameter concentrated deposits within that footprint, or alternatively with the entire area consisting entirely of massive sulfide. Individual vent fields can be separated by 10s to 100s of kilometers, depending on the geological setting (Hannington et al., 2011). Inactive sulfide-rich mineral deposits often surround active vents where mineral deposition is occurring, in a formation processes that can take thousands of years (Jamieson et al., 2013). The combined global footprint of all active vent ecosystems is estimated to be up to 50 km^2^, which is <0.00001% of the planet’s surface (Van Dover et al., 2018). Because of the difficulty in separating active from inactive vent sites (see Supplemental Materials), the total amount of mineable resources resulting from high temperature hydrothermal activity is covered in the next section on inactive sulfides.

From their first discovery in the late 1970s (Corliss et al., 1979), hydrothermal vents have attracted widespread public attention because they support unique and abundant animals that thrive in these systems because of symbioses with chemosynthetic microorganisms (Dubilier et al., 2008; Sievert and Vetriani, 2012). Geological and geochemical heterogeneity of vent fields leads to localized differences in fluid and deposit chemistry (Fouquet et al., 2010; German et al., 2016), which translates to animal endemism and biodiverse animal populations (Van Dover, 2000; Van Dover et al., 2018). Changes to hydrothermal venting chemistry or intensity could have repercussions on the types of microbial life that can exist, and therefore on the animals that can be supported.

#### 2.1.1. Possible impacts to biomass, primary production and microbial diversity at active vents

Microbes inhabit nearly every niche associated with active hydrothermal systems including the rocks and fluids in the subseafloor, sulfide chimney walls and surfaces, and in animal assemblages as internal and external symbionts (Fisher et al., 2007; Dubilier et al., 2008; Schrenk et al., 2009). Microbes growing on and within hydrothermal chimneys often produce thick biofilm mats easily visible to the naked eye, and even the extreme zones of high-temperature chimneys where temperatures up to 122°C host millions of microbial cells per gram of chimney material (Schrenk et al., 2003; Han et al., 2018). Nevertheless, the vast majority of microbial biomass in hydrothermal systems probably resides in the porous subseafloor underlying the chimneys. The amount of microbial biomass in the subseafloor of active hydrothermal vent systems is poorly constrained, though, due to difficulties in accessing this environment. Models of fluid circulation that assume a temperature limit of life of 122°C (Takai et al., 2008) as the main limitation to life yield a wide range of results depending on the depth of fluid circulation within the seafloor (Lowell et al., 2015), which can range from a few centimeters to the full thickness of the highly permeable basalt layer (~ 500 m). Therefore, mining activities risk removing the bulk of microbial biomass in areas where the habitable crustal area is thin. Without being able to directly observe and sample this microbial habitat, the diffuse fluids exiting cracks in the seafloor are considered to be windows into the subsurface of active vent systems (Deming and Baross, 1993; Huber and Holden, 2008). Diffuse fluids contain active microbial cells that are in one to two orders of magnitude greater abundance than that of the surrounding seawater (Karl et al., 1980; Huber et al., 2002; Meyer et al., 2013) with diverse metabolic capacities that affect global chemical cycling of carbon, nitrogen, iron, and sulfur (Mehta and Baross, 2006; Wankel et al., 2011; Holden et al., 2012; Bourbonnais et al., 2014; Fortunato et al., 2018).

The physiologically diverse microorganisms inhabiting active vent fields are considered to be fast growing and highly productive. The annual global production of biomass is estimated to reach 1.4 Tg carbon, significantly influencing deep-sea chemical cycling (McNichol et al., 2018). Thus, despite the small size of active vent fields, the rates of microbial primary productivity by microrganisms fueled by chemosynthesis in active vent systems can rival that of coastal and open ocean photosynthetic systems, making them productivity hotspots in an otherwise energy-starved deep sea (McNichol et al., 2018). As primary producers in the ecosystem, microbes support nearly all life at vents, from microbial consumers to the abundant and charismatic animals (Sievert and Vetriani, 2012). Animals can benefit from microbial primary productivity directly by harboring endosymbionts (e.g., tube worms, mussels, and clams) or indirectly by grazing microbial mats (e.g., *Rimicaris* shrimp)(Dubilier et al., 2008). Therefore, a major disruption of the chemical conditions that permit microbial chemosynthesis could have devastating consequences for all animals in that ecosystem.

Beyond their role in supporting primary productivity through chemosynthesis, many microbes in hydrothermal ecosystems play essential roles for animals by providing cues for larvae to settle (O’Brien et al., 2015). Microbial mats create barriers that slow the release of diffuse fluids and concentrate the energy-rich chemicals used in chemosynthesis. Growth of the microbial mat attracts microscopic and macroscopic grazers and traps more fluids, resulting in a rich ecosystem that supports dense and diverse assemblages of animals and microbes (Fisher et al., 2007). The concentration of energy and nutrients in complex, microbially-created habitats with strong spatial gradients fosters the evolution of highly diverse microbial communities, making hydrothermal vent systems hotspots of microbial diversity on the seafloor (Campbell et al., 2006; Schrenk et al., 2009; Olins et al., 2013; Meier et al., 2017).

In addition to the microscopic Bacteria and Archaea that are the base of the food web, hydrothermal systems also host abundant microscopic Eukarya, including protists and fungi (Edgcomb et al., 2002; López-Garcia et al., 2007), as well as viruses (Ortmann and Suttle, 2005; Williamson et al., 2008; Anderson et al., 2011; He et al., 2017). The diversity, distribution, and ecological roles of both groups are poorly constrained. Therefore, although hydrothermal systems are one of the better studied deep-sea environments (Figure 4), there is still much to learn about their microbial communities and their role in maintaining ecosystem functions.

**Figure 4.**
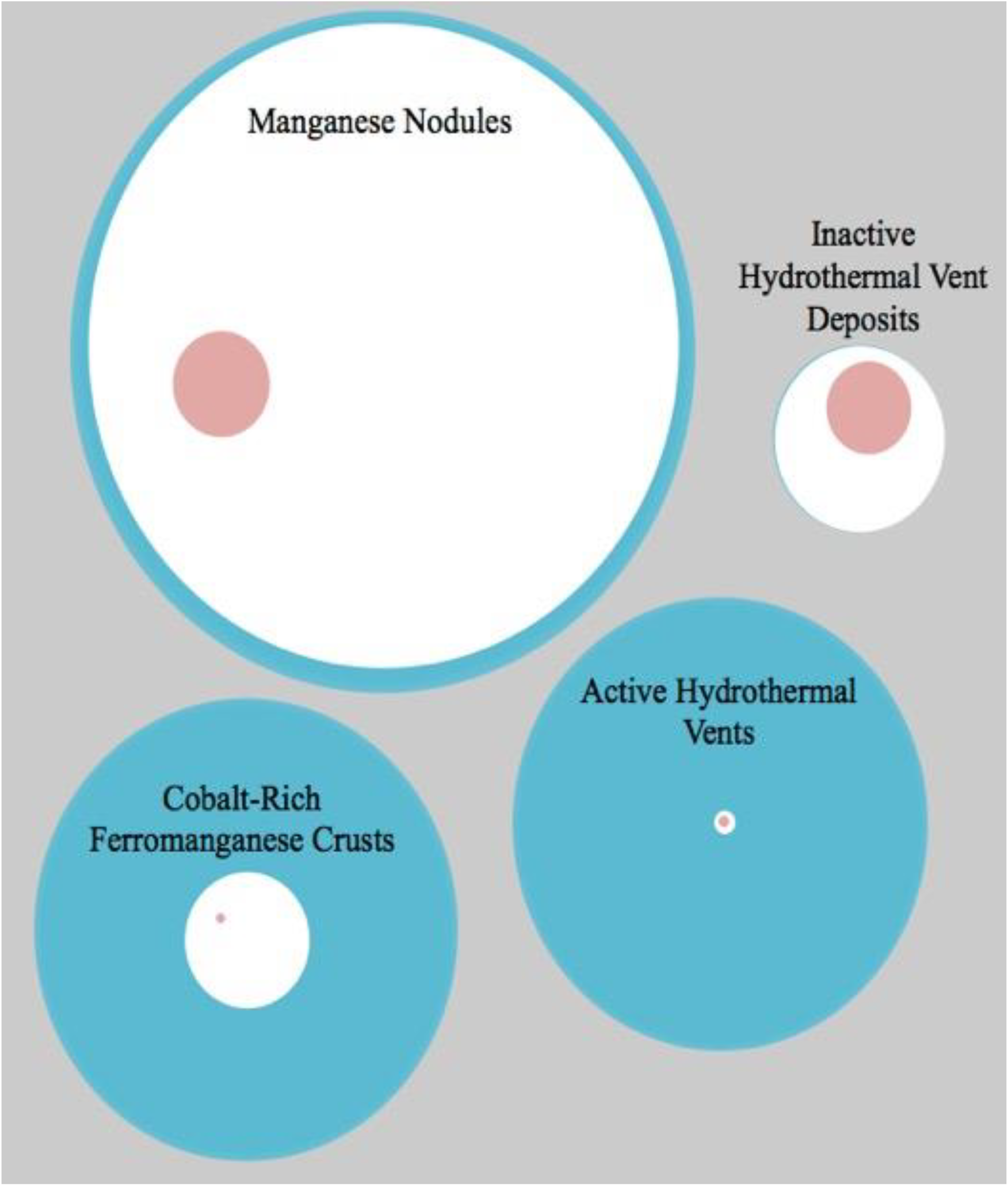
A schematic depicting the relative state of knowledge of deep-sea habitats with mineral resources versus the areal extent of the resources. White circles represent the relative area of the potentially exploitable resource (Petersen et al., 2016), and red circles signify the fraction of that area that is subject to current or pending exploration licenses (Hein et al., 2013a). Blue halos indicate the relative number of peer-reviewed publications with relevant key words. See Supplementary Materials Dataset 1 for more details.

In summary, microbial life at active hydrothermal vents are the dominant base of the food web at these sites, supporting abundant and diverse animal life at distinct “oases” on the seafloor. These microbial ecosystems comprise abundant standing stock of life that is diverse and highly productive, fueled by the abundant chemical energy supplies in these active vent systems. Mining activities near active vents could disrupt the nature of fluid flow, and therefore the availability of chemical energy to these ecosystems, potentially causing a cascade effect on the size, production, and diversity of these ecosystems (Figure 3) (Van Dover et al., 2018).

#### 2.1.2. Potential loss of genetic resources from active vent ecosystems

In addition to being biological hot spots, hydrothermal vents are also targets for natural products discovery owing to the unique genetic resources that some of the microbes contain (Thornburg et al., 2010). According to the Convention on Biological Diversity, the term “genetic resources” is defined as genetic material (i.e., any material of plant, animal, microbial, or other origin containing functional units of heredity) of actual or potential value. They may be used in biotechnological development of pharmaceutical drugs, research-based enzymes, food processing enzymes, cosmetic products, and other potential applications. Natural products from marine animals and microbes are already being marketed as anti-cancer and anti-viral drugs as well as various “cosmeceuticals” – cosmetic products with medicinal properties (Martins et al., 2014). For example, a novel benzoquinone compound isolated from a thermophilic bacterium from a deep-sea hydrothermal vent had anti-tumor activity by triggering cell death of cancer cells (Xu et al., 2017), and recently a novel antibiotic was identified from a vent microorganism (Shi et al., 2017). DNA polymerases Vent^®^ and Deep Vent®, both isolated from hyperthermophilic vent microorganisms, are marketed for research applications in molecular biology. Cosmeceuticals such as Abyssine^®^ and RefirMAR^®^ capitalize on excretions and internal proteins from hydrothermal vent bacteria and are marketed as reducing irritation in sensitive skin and reducing wrinkles. Numerous additional bioactive compounds from hydrothermal vent microorganisms are in research and development phases, and awareness of the potential of hydrothermal vents, other extreme environments, and the deep subseafloor in general, as sources of natural products is growing (Navarri et al., 2016; Zhang et al., 2018). The prevalence of symbiotic relationships among hydrothermal vent species may also be a source of untapped potential given that some bioactive compounds isolated from marine animals have now been attributed to their microbial symbionts (Piel, 2009; Penesyan et al., 2010).

Because each active vent site most likely contains endemic microbial species that are unique to the particular environmental conditions at that site (e.g. (Huber et al., 2010)), disruption of any vent site is likely to have some level of negative impacts on humanity’s ability to discover and utilize these genetic resources. The ability of these unique microbial ecosystems to reset after anthropogenic disturbance is poorly known, although there is evidence of recovery after natural disturbances such as volcanic eruptions (Opatkiewicz et al., 2009; Fortunato et al., 2018), so it is not clear if these impacts would be permanent, long-lasting, or ephemeral (Figure 3). Very few active vent systems have been studied in any detail, so a primary concern about mining of these systems is that their unique biodiversity and genetic resources could be lost before they are ever discovered.

#### 2.1.3 Impacts on other microbial ecosystem services at active vents

In addition to the metals that are sourced from hydrothermal vents, other reducing substrates such as methane – a potent greenhouse gas – are highly enriched at active vent sites (Holden et al., 2012). Some specialized microbes use methane as their primary energy source and convert it into the less potent greenhouse gas carbon dioxide. Microbial consumption of methane and other chemicals has a measurable impact on the flux of these chemicals from hydrothermal systems into the ocean (Wankel et al., 2011; Wankel et al., 2012), but more exploration of these systems is required for accurate estimates of their global contribution to the carbon cycle. Irrespective of its global significance, disruption of the vigorous microbial activity in hydrothermal systems is likely to have unpredictable consequences for nearby deep-sea habitats, which may be exposed to chemicals such as methane, hydrogen, and hydrogen sulfide that were previously removed in the vents (Figure 3).

Active vent systems also have incalculable value as representations of habitats that were likely to be prevalent on the ancient Earth and perhaps even acted as the cradle for the evolution of microbial life (Baross and Hoffman, 1985; Martin et al., 2008). Similar environmental conditions that promote vigorous microbial activity in vents today could have also promoted the origin and early evolution of life on ancient Earth (e.g. (Baaske et al., 2007)). Furthermore, the isolation of deep-sea vents from the surface would have enabled them to act as refugia for early lifeforms when conditions at the surface of the planet were not hospitable (Nisbet and Sleep, 2001). Many of the enzymes and metabolic pathways used by vent microbes today appear to contain clues about the nature of the first biological molecules (Russell and Martin, 2004) and key evolutionary milestones (Nasir et al., 2015). Therefore, the microbial diversity of active vents is not only important for modern ecosystem functions but also as natural wonders and precious cultural and educational resources that connect us to our ancient origins on this planet (Figure 3).

Finally, it must be emphasized that the vast majority of microbial life at hydrothermal vents has not been explored (Figure 4), despite increasing improvements in access to the deep ocean and new analytical tools (Xie et al., 2011; Fortunato and Huber, 2016; Fortunato et al., 2018). Therefore, many of the ecosystem services that microbes provide in these ecosystems are not yet known to science. Thus, the cultural heritage and educational services that active hydrothermal vents provide, both known and unknown, could be lost from mining activities.

In summary, the ecosystem services described above (Figure 3) highlight the importance of active hydrothermal vents. As argued elsewhere (Niner et al., 2018; Van Dover et al., 2018), given the substantial challenges associated with the meaning and measurement of “no net loss” guidelines – e.g., distinguishing between different types of biodiversity, the central importance of function in addition to identity, and consideration of spatial and time scales involved – we argue that active systems should not be mined. This view is bolstered by recent recommendations (IUCN, 2016; OECD, 2016; Cuyvers et al., 2018), which note that perturbation coupled with biodiversity offsets is not an acceptable interpretation of the ISA’s remit to protect the marine environment for the common benefit of humanity, and other recent studies that try to model the efficacy of offsets for this type of resource (Dunn et al., 2018). Caution is particularly prudent when there is uncertainty around an ecosystem’s vulnerability and recoverability (Donohue and al., 2016), as with microbial communities at active hydrothermal vents. Given this stance, it is troubling that a large fraction of known active vents are within areas of the seabed that have already been contracted for exploration and possible exploitation (Figures 3, 4).

### 2.2. INACTIVE VENT FIELDS WITH MASSIVE SULFIDE DEPOSITS

Inactive vent fields are the remnants of prior active hydrothermal circulation (Figure 2). Current volumetric estimates and ore percentages of seafloor massive sulfide deposits approximate those of terrestrial ores, though the size of individual deposits is up to 20 Mt as opposed to opposed to 50-60 Mt for terrestrial equivalents (Hoagland et al., 2010; Hannington et al., 2011; Petersen et al., 2016). 90% of known deposits are less than 2 Mt, with only 10% above the current 2 Mt threshold of economic interest (Petersen et al., 2016). Moreover, mineral content, and therefore economic value, also varies greatly between and within deposits, with those of largest volume not necessarily being the most valuable (Petersen et al., 2016). Finally, these are rough estimates and they do not distinguish between active and inactive hydrothermal vent fields, which could differ greatly in the challenges they present to mining operations and in the potential impacts to biological communities.

Inactive vent fields may be much more amenable to mining operations than active vents due to their size and absence of high-temperature acidic fluids. A complication, however, is that systems with no observable surficial venting may reveal underlying activity when disturbed by mining. Individual quiescent chimneys in a still-active vent fields are not a truly inactive hydrothermal system. Any indication of even minor venting of warm fluid could be indicative of high temperature fluids at depth, with any disruption of surface material from mining activities having operational and environmental consequences. Information is required regarding underlying hydrology and microbial colonization patterns before any predictions can be made regarding the potential unintended consequences of disturbing the microbial communities of inactive vent fields.

#### 2.2.1. Possible impacts to microbial biomass, primary production and diversity at inactive vent fields

Inactive vent fields are not currently known to host many endemic *animals*, though this may be due to a lack of exploration (Figure 4). Inactive vent fields represent a broad transition zone between actively venting hydrothermal systems and non-hydrothermal seafloor environments and are therefore expected to share features of each (Levin et al., 2016a; Levin et al., 2016b). As in active vents, many animal taxa in inactive vent fields obtain their nutrition in association with chemoautotrophic microbial symbionts (Erickson et al., 2009). Many animals from the next generation recruit their symbionts from the environment independently of the previous generation, so disruptions to the composition of the ambient seawater microbial communities could affect the ability of these animals to persist in areas adjacent to mining activities even if their own habitat is not directly affected (Figure 3).

By contrast, inactive hydrothermal vent fields are home to *microbial* species that are distinct from those of active hydrothermal sites (Suzuki et al., 2004; Erickson et al., 2009; Sylvan et al., 2012; Toner et al., 2013). The overall microbial community composition of inactive vent fields can be similar to that of the surrounding seafloor (Kato et al., 2010), indicating that inactive fields may not host as many unique and endemic populations as active vents do (Figure 3). Any generalized descriptions of inactive vent fields are premature, however, considering that very few examples have been detected and studied (Figure 4) (Boschen et al., 2013; Vare et al., 2018). Furthermore, very few studies have attempted to characterize the microbial communities of inactive vent fields, their roles in local and global biogeochemical cycling, or as refugia and seed-banks for the more dynamic active vent fields. Inactive hydrothermal systems may lack vigorous hydrothermal venting, but they nevertheless contain complex subsurface habitats with unknown microbial ecosystems. Ecosystem services that these subsurface microbial communities could potentially provide include primary production, secondary production, element cycling, and unique genetic resources, although knowledge of these services is poorly constrained due to very limited sampling (Figures 3, 4).

#### 2.2.2. Potential loss of habitat and creation of acidic conditions from mining massive sulfide deposits at inactive vent fields

A few categories of the potential impacts of mining on inactive sulfide-associated microbial communities are highlighted here. Our estimates mainly come from activities occurring within the national boundaries of Papua New Guinea, which is the most well-known mining project in a seafloor hydrothermal system, led by Nautilus Minerals Ltd. (Coffey Natural Systems, 2008). There have been recent reports of newer mining tests offshore Japan, but less information is publicly available from this site.

Mining seafloor massive sulfide deposits is a form of strip mining, where the top layer of sediment and crust is removed as overburden, and the exposed ore is removed in successive layers until the deposit is completely removed or “mined out”. By comparison to terrestrial mining sites, one can expect that exposure of massive sulfide deposits will start a cascade of abiotic and microbially catalyzed reactions, due to the exposure of the deposits to oxygenated seawater. Pyrite – an iron sulfide mineral – is the main constituent of massive sulfide deposits, and the overall oxidation reaction that occurs when this mineral is exposed to oxygen and water generates protons. Where the local environmental buffering capacity is unable to absorb these additional protons, a feedback system takes effect that causes a pH decrease. A change in pH causes changes in the type and speed of chemical reactions that occur (Bethke et al., 2011; Jin and Kirk, 2018), leading to changes in metal and oxygen dissolution properties in addition to changes in biology. This process and its effects are termed acid mine drainage or acid rock drainage in terrestrial systems (Schippers et al., 2010; Nordstrom, 2011), where many studies have been conducted on the pivotal roles that microbes play in contributing to these processes.

In terrestrial systems, exhausted strip mines create terraced open pits that can slowly fill with lakes or groundwater of altered chemistry, as any remaining metal-rich sulfides react with exposure to water and oxygen to create acidic conditions. In a marine sulfide system, such a pit will be permanently exposed to the oxic deep seawater long after extraction ceases, also allowing for the creation of acidic conditions. Although seawater has a higher pH buffering capacity than freshwater on land, a recent study on treatment of acid mine drainage from a terrestrial massive sulfide deposit (itself an ancient hydrothermal vent site) showed that a ratio of 1 part acid mine drainage to 90 parts seawater was required to neutralize the acid conditions (Sapsford et al., 2015). Biotic catalysis in the form of microbes may be a key factor in determining how exposed sulfide deposits will react to bottom seawater, but no studies have directly investigated the role of biological catalysis in marine environments affected by mining.

The consequences of the complete destruction and permanent loss of seafloor habitat caused by deep-sea mining are difficult to predict (Figure 3), since there is no precedent for such activities in the deep sea. One speculative scenario is that water in a deep mining pit on the seafloor may become sufficiently isolated from actively flowing seawater to stagnate and create a potentially permanent acidic, anoxic condition, but research is needed to investigate this possibility. Even minimal fluxes of material out of the pit could be sufficient to propagate acid mine drainage reactions to surrounding areas. Where there is local recharge of bottom seawater into ocean crust (Fisher and Wheat, 2010), which itself would likely be affected by changes in seafloor topography, the polluted water may also be entrained into the seafloor and transported from the point source farther than predicted. The degree to which acidic mining pits might influence surrounding ecosystems and the roles of microbial communities in these acid-generating reactions requires investigation.

#### 2.2.3. Generation of tailings plumes from mining massive sulfide deposits at inactive vent fields

In addition to the loss of habitat directly caused by mining the seafloor, mining activities will produce a plume of waste material that will disperse and fall on the surrounding seafloor, which is expected to nearly double the total area of seafloor impacted by mining (Boschen et al., 2013; Fallon et al., 2017). The environmental impact of this plume of waste material will depend on several factors, but perhaps most importantly, on the proximity of the waste plume to active hydrothermal systems. If the active vents are close enough to the mining area, they could become buried in the mining plume. Determination of this critical distance should be studied prior to any mining activities (Dunn et al., 2018).

Furthermore, because inactive vent fields are poorly explored and their possible level of hydrothermal activity is difficult to ascertain without high resolution seafloor surveys, there is a high likelihood that undiscovered active vents could be associated with apparently inactive fields. If these vents are not discovered prior to mining activities, they could become buried in the plume of waste material before there is any opportunity to explore their ecosystem services and potential scientific value. In addition, burial of seafloor habitat, even if it is not hydrothermally active, could disrupt the ability of animal larvae to sense seafloor conditions and to respond to environmental cues of where to attach and colonize (Gollner et al., 2010; Gollner et al., 2015).

There is also the potential impact of plumes of mining tailings closer to the ocean surface. Current mining operation designs propose to transport mined seafloor material to a surface ship for processing, returning the waste fluids to the ocean. This tailings waste stream, consisting of rock/ore fragments of small size and initial treatment chemicals as well as elevated concentrations of dissolved metals from the mining process, will create plumes of debris in the water column (Nath et al., 2012; Boschen et al., 2013; Fallon et al., 2017). Tailings may extend affected areas up to 80% from the point-source of pollution, though this can be difficult to predict, particularly in the deep sea where baseline information is scarce (Boschen et al., 2013; Hughes et al., 2015; Fallon et al., 2017; Vare et al., 2018). Studies on the effects and magnitude of potential metal leachate concentrations indicate significant local effects, particularly in areas of low or stagnant flow, and have the potential to remain in solution despite extensive mixing (Sapsford et al., 2015; Fallon et al., 2017; Fallon et al., 2018). Any substantial chemical amendments will likely have dramatic consequences for community structure and function, as observed with hydraulic fracturing on land (Murali Mohan et al., 2013).

Legislation currently requires treatment of any mine tailings on land to minimize this historically problematic waste (Dold, 2014; Hughes et al., 2015; Ma et al., 2017; Vare et al., 2018). Tailings often still contain elevated concentrations of acid-generating sulfides and heavy metals, and thus represent a significant additional source of mine drainage. Current deep-sea mining proposals indicate disposal of rock over the side of the mining vessel at various depths (Schriever and Thiel, 2013), which may eventually form deposits of sulfide-containing rock on the seafloor. This is similar to Deep-Sea Tailings Disposal, a strategy already used by a small number of terrestrial mines (Jones and Ellis, 1995; Schriever and Thiel, 2013; Dold, 2014; Vare et al., 2018). Measurable impacts from these tailings dumps include elevated concentrations of various transition metals in sediment, blanketing of the seabed by compacted precipitates, and release of elevated concentrations of sulfur and transition metals into the water column (Kline and Stekoll, 2001; Shimmield et al., 2007; Ramirez-Llodra et al., 2015; Hauton et al., 2017).

A few studies have investigated the various effects of these tailings plumes on animals (Kline and Stekoll, 2001; Mestre et al., 2017). Mining waste is known to affect microbial biogeochemical cycling and the rates and success of community recovery in shallow coastal sites (Pedersen, 1984; Pedersen and Losher, 1988; Almeida et al., 2007). However, no studies have explored the impacts of tailings plumes on deep-sea microbial communities. Natural plumes emitted from hydrothermal systems are known to have profound implications for the composition and activity of deep-sea microbial communities (Anantharaman et al., 2013; Dick et al., 2013; Levin et al., 2016a), therefore the potential impacts of tailings plumes can also be expected to be significant.

One concern is that disruption of natural microbial communities and stimulation of heavy metal-metabolizing microbes, in particular, will have far-reaching consequences for element cycling in the deep sea (Figure 3). Some metals may enter solution due to microbial activity, thus spreading the effect to a larger area and making the metals more bioavailable and increasing their toxicity. Others may precipitate out of solution more readily, causing issues such as blanketing areas of the seafloor with amorphous metal-rich precipitates. There is currently no research on the relevant thresholds over which some level of mining activity might begin to impact marine element cycling on a regional level.

Lessons from terrestrial massive sulfide mining show that environmental change brought about by these activities persists long after mining activity has ceased, including cases where point-source remediation measures are in place (Bird, 2016). However, remediation strategies that might be applied to operationally challenging deep-sea environments are poorly developed, though some studies have made tentative recommendations (Ramirez-Llodra et al., 2015; Vare et al., 2018). An industry report proposes that active sites will regenerate themselves by generating new mineral cover from already-present geochemical reactions and biology re-seeded from nearby refugia (Coffey Natural Systems, 2008), though the report does not specify how long this might take and whether it will require active human management. Even if this prediction is reasonable for active vent fields, it is not applicable to inactive vent fields. We speculate that taking no remedial action will likely result in acid mine drainage conditions over many decades, but research is need to assess this. Remediation strategies are likely to involve either natural dilution of the mining pit with seawater or else capping and permanent isolation of the pit. Both strategies have potential consequences and require extensive investigation. Depending on the local buffering capacities, natural dilution of the pit may not be sufficient to completely neutralize the acid-generating chemical reactions, potentially resulting in spreading acid-mine drainage across a much broader area of the seafloor. Capping the pit would require development of new technology capable of permanently isolating a large deep-sea pit, and failure of the cap could have devastating consequences for nearby ecosystems, potentially resulting in run-away acid mine drainage reactions within the capped region.

### 2.3. FERROMANGANESE NODULES

Ferromanganese nodules form in sediment underlying organic-poor regions of the global ocean, often at water depths > 4,000 meters (Figures 1, 2). In addition to iron and manganese, nodules incorporate high concentrations of economically valuable metals such as nickel, cobalt, and copper (Hein et al., 2013a). Nodule size ranges from microscopic particles to several cm in diameter and occur dispersed across nodule fields. Nodule growth is extremely slow (mm to cm accumulation per million years; (Ku and Broecker, 1965; Bender et al., 1966; Boltenkov, 2012)), and surrounding pelagic sediments accumulate Mn at approximately similar rates as nodules (< 5 mg cm^-3^ per 1000 years; (Bender et al., 1970)). Nodules can acquire manganese from sediment pore waters or from the overlying water column. The growth mechanism is mediated by the redox state of overlying waters and, in some environments, growth can be supported by hydrothermal influence and may change throughout the growth history of the manganese nodule (Mewes et al., 2014; Wegorzewski and Kuhn, 2014). Whether nodule growth proceeds purely abiotically, or is influenced by microbial activity or seeding is not currently known, although microbial communities have been detected in nodules (Tully and Heidelberg, 2013; Lindh et al., 2017). Recent studies have also documented novel animal communities that are supported by nodule fields (Bluhm et al., 1995; Purser et al., 2016; Vanreusel et al., 2016; Peukert et al., 2018).

Several studies have correlated water depth with nodule coverage to extrapolate and predict nodule occurrences over a wider area (Park et al., 1997; Jung et al., 2001; Kim et al., 2012; Peukert et al., 2018). Invariably, the occurrence of nodules coincides with areas of low sedimentation; for example, the sedimentation rate in the Clarion Clipperton Zone is estimated to be 0.3-15 mm per 1000 yrs (Jeong et al., 1994). These low sedimentation rates, which are typical of ocean gyres, are due to extremely low productivity of the overlying ocean, which exports low amounts of particles and organic matter to the deeper ocean. Thus, mining of nodules would disrupt deep-sea sediment environments that have evolved over millennia and would likely take just as long to recover to pre-disturbance conditions.

Manganese nodules harbor active microbial communities with cell densities three orders of magnitude higher than in surrounding sediment (Shiraishi et al., 2016). However, the specific organism(s) responsible for manganese oxidation and precipitation in those environments remain unidentified, despite some recent studies suggesting different chemical processes and structures occurring in the interiors versus exteriors of nodules, and nodule microbial communities that are distinct from the surrounding sediment (Tully and Heidelberg, 2013; Blöthe et al., 2015; Shulse et al., 2017). The interplay between sediment geochemistry and nodule microbial community structure remains poorly understood. It is therefore difficult to predict what the microbial and biogeochemical response and recovery would be to disturbance caused by deep sea mining (Figure 3).

#### 2.3.1. Limited impacts to organic carbon sequestration from mining ferromanganese nodules

Marine sediments are a major sink of organic matter over geological timescales and an important part of the global carbon and oxygen cycles (Berner, 2003). Sinking particles settling on the ocean floor are buried, effectively protecting and preserving their organic matter contents, and impeding it from microbial “remineralization” to carbon dioxide. The burial of organic carbon in the deep ocean is an important component of the global carbon cycle, thus regulating atmospheric CO2 and global climate through the sequestration of carbon, and allowing the build-up of oxygen in the atmosphere (Arndt et al., 2013; Hülse et al., 2017). Deep-sea mining of ferromanganese nodules will cause the re-suspension of sediments (Thiel and Schriever, 1990), potentially altering the ecosystem service of carbon sequestration that occurs in this habitat.

We estimate (see Supplemental Materials for calculations), however, that proposed mining of these nodule-bearing sediments and resulting re-suspension of particles and organic matter will have a trivial impact on the ecosystem service of carbon sequestration for two reasons (Figure 3). First, these sediments contain extremely low quantities of organic matter (<0.5% percent (Khripounoff et al., 2006). This is typical for deep-sea sediment (Seiter et al., 2004), since the particles delivering organic carbon to the ocean floor must sink over long distances to reach the ocean floor, during which the majority of organic matter is remineralized by microbes in the water column (Marsay et al., 2015; Cavan et al., 2017). Thus, only a relatively small mass of carbon might be re-suspended, compared to the much higher carbon loads in nearshore sediment environments. Second, the organic matter contained in these deep-sea sediments is likely to be highly processed and thus not particularly bioavailable to microbial remineralization, so most of the organic carbon would be redeposited on the seafloor and sequestered. Furthermore, as stimulation of organic carbon remineralization in the overlying water column is likely to be low, there would be inconsequential changes in dissolved oxygen concentration in bottom seawater (<0.5%).

#### 2.3.2. Other microbial ecosystem service impacts from mining ferromanganese nodules

Although the carbon sequestration ecosystem service of nodule fields would not be impacted, other microbial ecosystem services in nodule fields are expected to be impacted by mining activity (Figure 3). For example, as part of the European JPI Oceans Mining Impact project (Paul et al., 2018), the DISturbance and reCOLonization (DISCOL) area was recently revisited to study the long-term impact of nodule mining. The DISCOL experiment was carried out in 1989 in the Peru Basin in which the deep seafloor was plowed in an area of ~ 11 km^2^ to mimic nodule mining (Thiel et al., 2001). Clear geochemical differences, including metal distributions, in the upper 20 cm of disturbed and undisturbed sediments could be observed even 26 years after plowing (Paul et al., 2018). Based on their observations, the authors noted that nodule mining will likely have long-lasting impacts on the geochemistry of the underlying sediment (Paul et al., 2018). Specifically, solid-phase manganese concentrations were lower in disturbed areas compared to reference areas. This finding suggests that the capacity for metal sequestration via scavenging onto nodules will be substantially limited during the recovery period. The absence of nodules in the disturbed area increases metal flux out of sediment, although it is argued that these flux rates do not reach rates that are potentially toxic to animals (Paul et al., 2018).

Nodule regrowth may also be limited by both the geochemical and microbiological changes following mining-related disturbances. For example, thermodynamic and kinetic constraints limit the oxidation of reduced manganese to oxidized manganese by oxygen (Luther, 2010). Microbes can catalyze this reaction via direct and indirect pathways; thus, the formation of most manganese oxide minerals in the environment is microbially mediated (Hansel and Learman, 2015). A broad diversity of organisms are capable of manganese oxidation, from bacteria to fungi (Hansel, 2017), although microbial manganese oxidation does not provide an energetic benefit to the organism and the physiological purpose is unclear.

Mining activities will cause a decrease in the ecosystem service of this habitat through the destruction of paleoscientific records, a valuable education aspect of this environment. Marine sediment cores are an immensely valuable resource for reconstructing climate conditions over Earth’s history (Figure 3). Plant and animal fossils found in sediments are frequently used to reconstruct and understand the past chemistry and temperature of the ocean. For example, the calcium carbonate shells of microorganisms such as foraminifera or coccoliths can be analyzed using oxygen isotopes to determine the temperature and chemistry of ancient seawater and how cold the ocean was at the time the shell formed (Spero et al., 1997; Ornella Amore et al., 2004; Maeda et al., 2017). Moreover, diatom microfossils can be used to understand upwelling currents and reconstruct past wind and weather patterns (Abrantes, 1991; Schrader and Sorknes, 1991; Zúñiga et al., 2017). In addition, dust layers found in sediment cores can be analyzed to determine its origin to understand the direction and strength of winds and how dry the climate may have been at that particular time (Rea, 1994; Middleton et al., 2018). Nodules themselves record paleoclimate and seawater conditions. For example, rare earth element ratios in nodules serve as a proxy for bottom water redox state and changes in deep currents over time (Glasby et al., 1987; Kasten et al., 1998), and some rare earth element isotopes in ferromanganese nodules also reveal patterns in ocean circulation throughout time (Albarède et al., 1997; Frank et al., 1999; van de Flierdt et al., 2004). Marine sediment and nodules thus serve as a valuable resource for reconstructing past climate conditions as well as understanding and predicting future climate change. Sediment in nodule-rich regions is particularly valuable due to a low sedimentation rate that allows for piston coring techniques to readily access extremely old sediment. Widespread sediment disturbances from nodule mining would result in the loss of this record and educational ecosystem service (Figure 3).

Overall, mining activities in nodule fields will have varied impacts on microbial ecosystem services (Figure 3). Some services, such as carbon sequestration potential, will be minimally impacted. Other services, such as research and educational value from paleoscientific records contained with sediment layers, would be severely perturbed and not recoverable. Decades-long studies have identified that microbial processes with the sediments underlying nodules remain impacted for quite some time (Paul et al., 2018), but the corresponding impact this has to biogeochemical cycling and ecological functioning is not constrained and requires further investigation, despite this resource type having been the most studied for these kinds of impacts (Figure 4).

### 2.4. COBALT CRUSTS ON BASALTIC SEAMOUNTS

Cobalt-rich crusts (also called polymetallic crusts) occur on sediment-free rock surfaces in all oceans of the world (Figures 1, 2), raging in thickness from <1 mm to ~260 mm. They are most common in the Pacific Ocean where there are estimated to be over 50,000 seamounts and knolls and many more seamounts likely exist in uncharted waters (Wessel et al., 2010; Levin et al., 2016c). In addition to cobalt, other rare and trace metals of high economic value – including copper, nickel, platinum and tellurium (used in the solar cell industry) – are adsorbed to the crust from seawater. In the central Pacific, ~7,500 million dry tons of crusts are estimated, containing 4 times more cobalt, 9 times more tellurium, and a third of the manganese that makes up the entire land based reserve of these metals (Hein et al., 2013b). Polymetallic crusts are formed slowly (1-5 mm per million years), and biomineralization by microorganisms plays a role in initiation of crust accretion, serving as a biological nuclei (Wang and Müller, 2009). Microorganisms may also play a role promoting the enrichment of cobalt in the crust through sorption/immobilization processes (Krishnan et al., 2006; Sujith et al., 2017).

#### 2.4.1. Possible impacts to biomass, primary production and microbial diversity in cobalt crusts

The alteration rinds that form on seafloor exposed basalts at seamounts and outcrops (i.e. cobalt-rich crusts) provide a habitat suitable for sessile animals like corals and sponges that require a hard substrate to attach to (Etnoyer et al., 2010; Shank, 2010), as well as for brooding animals like octopus (Hartwell et al., 2018). However, the role microorganisms play in faunal colonization and presence in these regions remains unknown, as does the relative role of microbial chemosynthesis and heterotrophy in this ecosystem. Some studies suggest persistent patterns in microbial community composition on highly altered seafloor-exposed basalts that have ferromanganese crusts (Lee et al., 2015). Surveys of indigenous microorganisms from sediments associated with cobalt-rich crusts (Liao et al., 2011; Huo et al., 2015) and manganese rich crust (Nitahara et al., 2011) have detected the potential for microbial chemosynthetic primary production supported by ammonia oxidation. Similarly, the amount of primary production supported by microbial communities on altered seafloor basalts could be significant for carbon cycling in the deep sea (Orcutt 2015).

Removal of alteration crusts from seamounts and outcrops through mining/dredging is expected to physically alter the seafloor substantially. The overall slope of the seamount may be flattened, and the amount of soft sediment increased though disturbance and release of waste during the mining process (Levin, 2013). This mining activity would dramatically impact sessile animal communities, although recovery rates from these disturbances are unknown. Mining activity would expose fresh surfaces of underlying basalt rocks, which would eventually be altered through seawater exposure, although this process would be very slow (1-5 mm per million years). Due to the slow growth of both the alteration crust as well as the fauna that live on them, the recovery time for physically disturbed crusts on seamounts caused by mining is predicted to be long (Schlacher et al., 2014). For example, recovery and recolonization of seamounts by animals was found to not have occurred after 10 years of closing bottom-trawling activities in coastal New Zealand and Australia (Williams et al., 2010). Dredging activities for recovering the crusted rocks underneath, and not just animals, is expected to have even slower recovery rates.

Moreover, mining/dredging activities that change the physical structure of the seamount/outcrop would potentially impact fluid circulation pathways through basaltic crust, especially on ridge flanks. This outcrop-to-outcrop fluid circulation away from the ridge axis ventilates the majority of heat from the oceans (Fisher et al., 2003; Fisher and Wheat, 2010) and is an important component of global geochemical cycles (Wheat et al., 2017; Wheat et al., submitted). For example, a recent study indicated that at least 5% of the global ocean dissolved organic pool is removed via microbial oxidation within the subsurface ocean crust during ridge flank fluid circulation (Shah Walter et al., 2018). Mining activities could change the permeability, porosity and locations of fluid discharge, which would impact fluid circulation and could have consequences on the nature of microbial communities resident in these environments that are influenced by fluid conditions (Figure 3) (Zinke et al., 2018). However, very little is known about fluid circulation far away from ridge axes where many seamounts occur, so it is difficult to know how widespread this disruption could be.

#### 2.4.2. Other possible impacts to microbial ecosystem services in cobalt crusts

Microorganisms in cobalt crusts can likely use metals as an energy source and carry adaptations to tolerate the high heavy metal concentrations that occur in polymetallic crusts, potentially playing a role in metal cycling in oceans. Crustal microorganisms have demonstrated the ability to immobilize cobalt from seawater, release trace metals like nickel, and may also be capable of scavenging other metals (Krishnan et al., 2006; Antony et al., 2011). These traits are of interest for biotechnological applications, or applications that involve metal/microbe interactions such as bioremediation of polluted sites, bioleaching, and metal recovery (Figure 3). However, the financial considerations for dredging cobalt crusts from the seafloor limit the viability of this natural product discovery track.

## 3. RECOMMENDATIONS FOR BASELINE AND MONITORING DATA TO EVALUATE IMPACTS TO MICROBIAL ECOSYSTEM SERVICES

Recommendations for baseline measurements and monitoring of mining impacts have been published elsewhere (Gjerde et al., 2016; Henocque, 2017; Boetius and Haeckel, 2018; Cuyvers et al., 2018; Durden et al., 2018; Jones et al., 2018). These recommendations include measurements of animal biodiversity and deep-sea ecosystem structure and function. We propose that including microorganisms in these biodiversity and ecosystem measurements is critical for effective monitoring of mining impacts. In high energy environments like hydrothermal vents, where many microorganisms have short generation times, measurements of microbial diversity are likely to be highly sensitive and quickly responsive to environmental impacts. Alternatively, impacts in lower energy environments like deep-sea sediments hosting ferromanganese nodules may be harder to discern. Furthermore, the responses of microbial communities to mining impacts will be more complex than a simple "good" or "bad", as microbial species will respond in many different ways. Changes in microbial community composition are likely to convey a wealth of information about changes to the environment, if we are able to detect and decipher these signals. With enough research, monitoring of microbial communities could become sufficiently sensitive and specific to enable adjustments of ongoing mining activities before impacts to animal communities reach a dangerous threshold.

Predicting and assessing the environmental impacts of mining in the deep sea is fundamentally more challenging than on land because so little of the deep sea has been explored in any detail. In many areas under consideration for mining, we lack any knowledge of how the resident microbial communities contribute to primary production and element cycling in their habitats and how these local activities relate to regional- and global-scale chemical cycles. Therefore, any assessment or monitoring of mining impacts should consider the potential unexpected consequences associated with undiscovered microbial organisms and activities stimulated, directly or indirectly, by mining activities.

One pragmatic approach to the monitoring of mining impacts is the creation of protected areas and reserves, as recommended by others. Protected areas, such as Preservation Reference Zones where no impacts occur within mining sites (International Seabed Authority, 2018), and reserves would be particularly useful as reference points for the monitoring of microbial communities, since there is no way to assess the status of a microbial community *a priori* without reference points. Samples for detailed microbial community analysis through DNA sequencing approaches should be collected from both protected and impacted sites, to evaluate change. It is worth noting, though, that the technology for measuring microbial diversity is advancing so quickly that baseline measurements collected prior to mining activities are likely to be rendered obsolete shortly thereafter, therefore appropriate samples should be archived for re-analysis with new techniques as they become available. We also recommend the use of in situ and lab-based activity-oriented experiments to evaluate changes in metabolic activities that could alter element and nutrient distributions due to anticipated disruptions.

In conclusion, while some ecosystem services provided by microbial life in deep-sea habitats may be minimally impacted by mining activities, others are expected to be severely impacted (Figure 3). Active vent environments are expected to suffer the most extreme impacts from mining activity, which will be hard to avoid even with protected offsets (Dunn et al., 2018). There are several critical knowledge gaps that remain, and these are not evenly distributed across habitat type (Figure 4). For example, the long-term impacts of mining inactive sulfide deposits are poorly known, but could be dramatic in comparison to open-pit mines on land. When considering that the total estimated copper and zinc potential of these deposits are only slightly larger than the annual production on land (Hannington et al., 2011), it is important to weigh the consequences of these activities in environmental impact assessments. Moreover, it is unclear how extensively the seabed and overlying water column can be disturbed before tipping points are reached and some ecosystem services become negatively and/or critically impacted on local and regional scales. We highly recommend that baseline assessments of microbial diversity, biomass, and rates of hemical processes be included in environmental impact assessment planning, as they are currently lacking in policy recommendations (International Seabed Authority, 2012).

## ACKNOWLEDGEMENTS

We thank the other participants in the April 2018 workshop at Bigelow Laboratory for Ocean Sciences for their contributions to the ideas summarized in this manuscript: David Emerson, Kristina Gjerde, Susan Lang, Jennifer Le, James J. McManus, Ramunas Stepanaukas, Jason Sylvan, Margaret K. Tivey, and Geoff Wheat. We also thank Verena Tunnicliff and Cindy Lee Van Dover for providing advice on this topic, Kim Juniper for providing input on the manuscript, James Hein for providing the shape file information used in Figure 1, and Brandy Toner and Cara Santelli for providing photomicrographs used in Figure 2. This is C-DEBI publication number XXX.

## FUNDING STATEMENT

The Center for Dark Energy Biosphere Investigations (C-DEBI, funded by the US National Science Foundation award OIA-0939564) and the Deep Carbon Observatory at the Carnegie Institution of Washington (funded by the Alfred P. Sloan Foundation, CIW subaward 10693-03) are gratefully acknowledged for their funding support for the workshop and development and publication of this manuscript.

## AUTHOR CONTRIBUTIONS STATEMENT

RMJ constructed the maps in Figure 1, JJM developed the dataset for Figure 5, JAH contributed to Figure 2, and BNO created the remaining figures. BNO wrote the manuscript with input from all authors.

## CONFLICT OF INTEREST STATEMENT

The authors declare no conflicts of interest.

